# SirenScan reveals the conserved presence of siren RNAs and their evolutionary diversification across angiosperms

**DOI:** 10.64898/2026.08.14.744852

**Authors:** Haoran Peng, Aurélien Valentin, Yichun Qiu, Katarzyna Dziasek, Claudia Köhler

**Affiliations:** Department of Plant Reproductive Biology and Epigenetics, Max Planck Institute of Molecular Plant Physiology, Potsdam, Germany; Department of Plant Biology, Uppsala BioCenter, Swedish University of Agricultural Sciences and Linnean Centre for Plant Biology, Uppsala, Sweden

## Abstract

- Siren RNAs are exceptionally abundant reproductive small interfering RNAs (siRNAs), yet whether they represent a conserved feature of angiosperm reproduction has remained unclear because siren loci lack a standardized definition and identification strategy.
- We developed SirenScan, a computational framework that integrates cumulation-based and density-based analyses to identify siren loci from small RNA sequencing data. Using SirenScan, we systematically compared ovule and vegetative tissues across representative angiosperm lineages, including the early-diverging angiosperm *Nymphaea colorata*.
- We demonstrate that the presence of siren RNAs is a conserved feature of angiosperm ovules, although the degree of expression dominance varies substantially among species. Siren loci are consistently enriched within transposable elements (TEs), while the associated TE superfamilies and *cis*-regulatory motifs exhibit lineage-specific diversification. In Brassicaceae, siren loci share conserved CLASSY3-associated sequence motifs, whereas other angiosperm lineages display distinct motif compositions, suggesting evolutionary diversification of Polymerase IV targeting mechanisms.
- Our findings establish siren RNAs as a conserved component of the angiosperm reproductive small RNA landscape and support a model in which conserved RNA-directed DNA methylation machinery is coupled with lineage-specific regulatory mechanisms and transposon landscapes to shape siren locus evolution. SirenScan provides a robust and standardized framework for comparative studies of siren RNA biology and reproductive epigenomics across plant species.

## Introduction

Small interfering RNAs (siRNAs) are central components of RNA silencing pathways in plants, playing crucial roles in transcriptional and post-transcriptional gene regulation, genome defense, and epigenetic inheritance (Vaucheret & Voinnet, 2024). Among these, 24-nucleotide (nt) siRNAs are central to RNA-directed DNA methylation (RdDM), a key epigenetic mechanism that establishes and maintains DNA methylation, particularly at transposable elements (TEs) and other repetitive sequences (Matzke & Mosher, 2014; Cuerda-Gil & Slotkin, 2016). The canonical RdDM pathway depends on a plant-specific transcriptional machinery centered on RNA polymerase IV (Pol IV) (Herr *et al*., 2005) and RNA polymerase V (Pol V) (Wierzbicki *et al*., 2008), together with RNA-DEPENDENT RNA POLYMERASE 2 (RDR2) (Xie *et al*., 2004), DICER-LIKE 3 (DCL3) (Xie *et al*., 2004), and ARGONAUTE4 (AGO4) (Zilberman *et al*., 2003), which collectively generate and utilize 24-nt siRNAs to guide DNA methylation. While the core RdDM machinery is conserved across land plants (Trujillo *et al*., 2018; Bélanger *et al*., 2023), RNA Pol IV targeting factors have diversified in a lineage-specific manner, enabling flexible chromatin-directed deployment of siRNA–mediated silencing (Trujillo *et al*., 2018; Chakraborty *et al*., 2024).

Plant reproductive tissues exhibit highly distinctive small RNA (sRNA) landscapes compared with vegetative tissues (Petrella *et al*., 2021). Early studies in *Arabidopsis thaliana* revealed that developing endosperm accumulates abundant Pol IV-dependent 24-nt siRNAs that are predominantly maternal in origin (Mosher *et al*., 2009). Subsequent work demonstrated that these maternal siRNAs are essential for proper endosperm development, allelic dosage balance, and regulation of imprinted gene expression (Lu *et al*., 2012; Erdmann *et al*., 2017; Kirkbride *et al*., 2019). While characterizing the siRNA landscape of rice (*Oryza sativa* ssp. *japonica*) endosperm, it was observed that a very small number of genomic loci produces a disproportionate fraction of total siRNA molecules. These loci were initially termed <u>s</u>mall-interfering <u>R</u>NAs in <u>en</u>dosperm (siren) loci (Rodrigues *et al*., 2013). Following studies demonstrated that highly expressed siren RNAs are not restricted to rice endosperm but are also present in female reproductive tissues of other species, including *Arabidopsis thaliana* (Grover *et al*., 2020; Zhou *et al*., 2022; Burgess *et al*., 2022), *Brassica rapa* (Grover *et al*., 2020; Burgess *et al*., 2022), and *Capsella rubella* (Dziasek *et al*., 2024). Notably, siren RNAs were detected not only in the endosperm but also in ovules prior to fertilization and in the developing seed coat after fertilization (Rodrigues *et al*., 2013, 2021; Grover *et al*., 2020; Zhou *et al*., 2022; Li *et al*., 2022; Burgess *et al*., 2022; Dziasek *et al*., 2024). Despite this broader tissue distribution, the term “siren” was retained, reflecting both their original discovery context and their defining property: a small subset of loci that overwhelmingly dominate the siRNA landscape, metaphorically acting as “loud” loci that surpass all others in siRNA output (Grover *et al*., 2020). Functional studies in *Brassica rapa* showed that siren loci constitute the dominant source of maternal siRNAs required for normal seed development (Grover *et al*., 2018, 2020). Moreover, in *Capsella rubella*, dosage-sensitive maternal siren siRNAs were demonstrated to determine hybridization success and seed viability, directly linking siren loci to parental genome balance and postzygotic reproductive barriers (Dziasek *et al*., 2024). Together, current findings converge on a model in which siren RNAs are predominantly produced in ovules and subsequently influence epigenetic regulation in seed tissues, although the precise spatial dynamics of siren RNA production and action remain to be resolved.

Siren RNA biogenesis relies on locus-specific targeting of Pol IV, which is mediated by transcription factors including members of the reproductive meristem (REM) family and GENETICS DETERMINES EPIGENETICS1 (GDE1) (Wu *et al*., 2025; Xu *et al*., 2025; Pandesha & Slotkin, 2025). This transcriptional guidance operates in concert with CLASSY (CLSY) proteins, putative chromatin remodeling factors that stabilize Pol IV at selected loci (Felgines *et al*., 2024). In *Arabidopsis*, the CLSY3 paralog displays specificity for ovules and tapetal tissues and preferentially regulates siren loci in these reproductive tissues (Zhou *et al*., 2022, 2018). The CLSY homologs in rice similarly control reproductive siren RNA landscapes and genomic imprinting (Xu *et al*., 2024; Pal *et al*., 2024, 2025; Zhang *et al*., 2026). Together, these findings support a conserved regulatory mechanism in angiosperms, in which transcriptional and chromatin-based cues are integrated through CLSY-mediated Pol IV targeting to enable tissue-specific siren RNA production.

Despite increasing mechanistic insight into the regulation and function of siren RNAs, their identification from sRNA sequencing data remains methodologically challenging. Defining an objective boundary between siren and non-siren loci has proven to be non-trivial. A first attempt was made in rice endosperm, which examined the distribution of siRNA-producing loci using expression density plots based on reads per kilobase million (RPKM) values. The results showed a pronounced right-skewed tail, with a small number of loci exhibiting expression levels orders of magnitude higher than the genome-wide mean. However, no statistical threshold was proposed to distinguish siren loci from the broader population of siRNA-producing loci (Rodrigues *et al*., 2013). Following this initial observation, no dedicated methodological framework for siren locus identification was developed until a recent study introduced a cumulation-based strategy (Grover *et al*., 2020). In this approach, siRNA-producing clusters were ranked by expression level based on reads per million (RPM), and the minimal set of loci required to account for a specified fraction of total siRNA abundance— typically 90%—was designated as siren loci. Variants of this cumulation-based method have since been adopted in multiple studies (Grover *et al*., 2020; Zhou *et al*., 2022; Li *et al*., 2022; Pal *et al*., 2024; Dziasek *et al*., 2024; Kim *et al*., 2024). Despite their widespread use, they rely on user-defined cutoffs that lack a unified statistical rationale. In practice, the cumulative abundance threshold has been adjusted across studies to accommodate different datasets, ranging from 60% to 90% (**Table 1**), while density-based approaches similarly depend on visually inferred boundaries. Furthermore, the analytical pipelines used to generate siRNA clusters vary substantially among studies, including differences in read processing, clustering algorithms, and parameter settings (**Table 1**). These methodological inconsistencies complicate direct comparisons of siren loci across tissues, species, and publications.

**Table 1.**
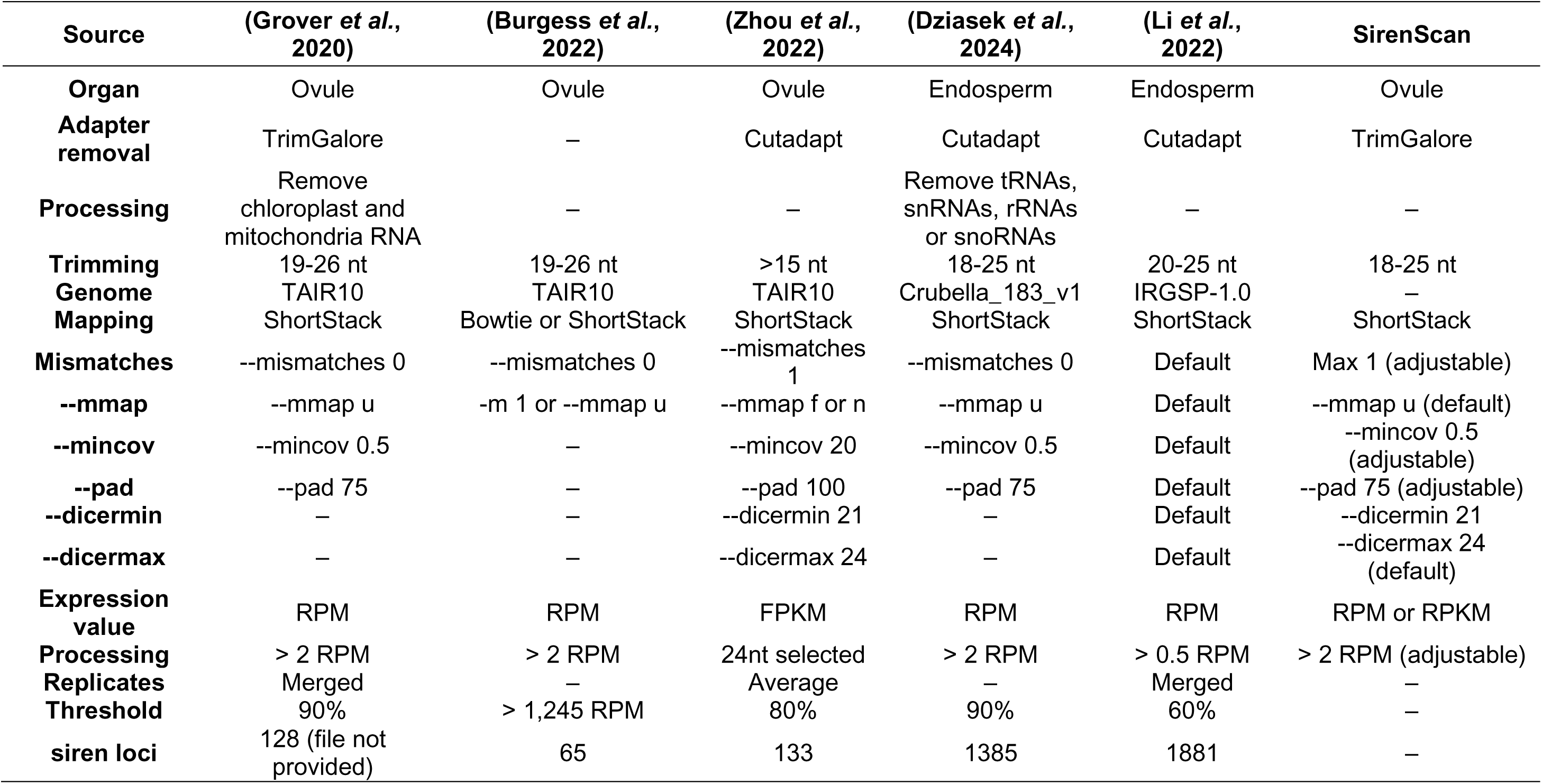
Summary of computational parameters and pipelines used for siren loci identification.

Here, we introduce SirenScan, a computational pipeline for identifying siren RNAs by integrating rank-abundance and density-based analyses of sRNA sequencing data. Applying SirenScan to ovule and leaf tissues from five angiosperm species, we aimed to systematically assess the conservation of siren RNA presence across plant lineages and to disentangle lineage-specific patterns of siren locus diversification. Using this unified analytical framework, we further examined the relationship between siren loci and TE composition, as well as the contribution of *cis*-regulatory features to siren locus specification. Together, this study establishes SirenScan as a robust and generalizable pipeline for comparative analysis of siren RNAs and provide new insights into the evolutionary and regulatory principles shaping extreme siRNA expression dominance in plant reproduction.

## Materials and Methods

### Plant material and growth conditions

*Capsella rubella* accession *Cr1GR1* was grown in a controlled growth chamber under 60% relative humidity with a 16 h light / 8 h dark photoperiod (21 °C day / 18 °C night) and a light intensity of 150 μmol photons m⁻² s⁻¹. *Solanum lycopersicum* cv. *Moneymaker* plants were grown under standard greenhouse conditions with a light intensity of approximately 250 μmol photons m⁻² s⁻¹. *Nymphaea colorata* (Casp.) Verdc. plants were cultivated under standard aquatic growth conditions. Leaf tissues from adult plants and mature ovule samples were collected from independent individuals and immediately frozen in liquid nitrogen prior to RNA extraction.

### RNA extraction and small RNA sequencing

Total RNA was extracted using the Spectrum™ Plant Total RNA Kit (Sigma-Aldrich, STRN50) following the manufacturer’s instructions. Protocol A was used to ensure recovery of small-sized RNA molecules. Small RNA library preparation and sequencing were performed by Biomarker Technologies (BMK) GmbH. Libraries were constructed using the VAHTS Small RNA Library Prep Kit for Illumina (Vazyme, NR801) according to the manufacturer’s protocol. Briefly, 3′ and 5′ adapters were sequentially ligated to small RNAs, followed by reverse transcription and PCR amplification. Amplified products were size-selected by PAGE gel electrophoresis and libraries were purified and quality-controlled prior to sequencing. Qualified libraries were sequenced on the Illumina NovaSeq platform.

### SirenScan pipeline for small RNA sequencing analysis

Small RNA sequencing data were processed using the SirenScan pipeline. Raw FASTQ files were used as input unless otherwise specified. Initial quality control of sequencing reads was performed using FastQC (Andrews, 2010). Adapter sequences were identified for each library using FindAdapt (Chen *et al*., 2024), based on the specified organism, unless adapter sequences were explicitly provided by the user. Adapter trimming was carried out using Trim Galore (Krueger, 2015) in combination with Cutadapt (Martin, 2011), retaining reads between 18 and 25 nt in length. For datasets that had been preprocessed externally, trimming could be skipped. A second round of quality control was performed by visualizing read length distributions to verify trimming efficiency. Trimmed reads were mapped to a reference genome using ShortStack v4.0.3 (Axtell, 2013), with alignments generated by Bowtie v1.3.1 (Langmead *et al*., 2009). Mapping parameters, including the number of allowed mismatches, were adjustable. Alternatively, pre-aligned BAM files could be provided as input. siRNA producing loci were defined by ShortStack clustering, with parameters controlling cluster merging distance and minimum coverage configurable by the user. Following clustering, expression levels were quantified either as reads per million mapped reads (RPM) or reads per kilobase per million mapped reads (RPKM). Analyses could optionally be restricted to uniquely mapped reads to minimize contributions from highly repetitive sequences. For each sample, siren loci were identified using two complementary strategies. In the cumulation-based approach, siRNA clusters were ranked by expression level, and the minimal set of loci required to account for a user-defined fraction of total siRNA abundance was selected. In the density-based approach, the distribution of cluster expression values was analyzed to identify highly expressed loci. When expression distributions exhibited bimodality, siren loci were defined as clusters belonging to the high-expression mode separated by a local minimum. In cases where distributions showed a long right-skewed tail without clear bimodality, siren loci were defined as clusters with expression levels exceeding a specified multiple of the median expression. Details of the small RNA sequencing datasets and genome information (Lamesch *et al*., 2012; Slotte *et al*., 2013; Zhang *et al*., 2020a; Shang *et al*., 2023; Chen *et al*., 2026) for each species are provided in **Supplementary Table 1**. Parameters used for siren locus identification in each species are summarized in **Supplementary Table 2**.

### Motif enrichment analysis

Motif enrichment analysis of siren loci was performed using MEME (v5.5.9) in classic mode (Bailey *et al*., 2015). DNA sequences corresponding to siren loci were analyzed with the following parameters: zero or one occurrence per sequence (ZOOPS), identification of up to three motifs, and motif widths ranging from 6 to 50 bp. Both strands were considered during motif discovery.

### Permutation test for enrichment analysis

Similarly to a previous study (Batista *et al*., 2019), the enrichment of siren loci within genomic features was assessed using permutation tests implemented in the R package regioneR v1.8.1 (Gel *et al*., 2016) implemented in R version 4.4.1. Siren loci and genomic annotations, including genes and TE superfamilies, were represented as genomic intervals. For each feature, the observed number of overlaps with siren loci was calculated and compared to a null distribution generated from 10,000 random permutations. In each iteration, siren loci were randomly redistributed across the genome using the randomizeRegions function while preserving chromosome identity and region length. Overlaps were recalculated using the numOverlaps function. Empirical p-values and Z-scores were computed based on the distribution of randomized overlaps. To control for the strong genomic bias imposed by LTR/Gypsy rich heterochromatin in *Solanum*, permutation tests were repeated using a masked genome excluding LTR/Gypsy regions.

### CLSY3/4 sequence alignments

AtCLSY3/4 protein sequences were retrieved from TAIR10 (https://www.arabidopsis.org) and used in BLASTP searches to identify CLSY3/4 proteins in the other investigated species **(Supplementary Table 3)** based on reciprocal best BLAST hits. MUSCLE (Edgar, 2004) was applied with default settings to generate multiple sequence alignments.

## Results

### Overview of the SirenScan analytical workflow

SirenScan is a flexible and integrative pipeline designed to identify siren loci as dominant contributors to sRNA populations by combining cumulation-based and density-based analyses within a single workflow **(Fig.1)**. Starting from sRNA sequencing data, SirenScan performs standardized preprocessing, including automated adapter detection, read trimming, and quality control, followed by genome alignment and clustering of sRNA-producing loci by ShortStack (Axtell, 2013). Reads are mapped to a reference genome and grouped into clusters representing discrete siRNA-producing regions, from which normalized expression metrics such as RPM or RPKM are derived. The core analytical step of SirenScan is the identification of siren loci using two complementary strategies. In the cumulation-based approach, siRNA-producing loci are ranked according to expression level, and cumulative abundance curves are computed to determine the minimal set of loci required to account for a user-defined fraction of total siRNA output. This strategy directly captures the defining feature of siren RNAs—expression dominance—and enables quantitative comparison of dominance strength across tissues and species. In parallel, SirenScan applies a density-based strategy that models the distribution of cluster expression values. Highly expressed loci are identified either by detecting a local minimum separating high– and low-expression populations in bimodal distributions or, when such separation is absent, by applying a median-based cutoff that identifies extreme outliers in right-skewed distributions. By integrating these strategies, SirenScan avoids reliance on a single arbitrary threshold and provides internal validation of siren locus identification. Beyond locus identification, SirenScan generates a comprehensive suite of diagnostic and comparative outputs that facilitate interpretation of siRNA expression architectures **(Supplementary Fig.1a-h)**. These include cumulative rank– abundance plots, expression density profiles, and scatter plots that visualize individual clusters within the global expression landscape. Color-consistent scatter plots enable tracking of the same loci across multiple samples, while Venn diagrams and UpSet plots summarize overlap and specificity of siren loci between tissues. Together, these outputs allow users to assess both the presence and the relative strength of siren RNA dominance across datasets.

**Fig. 1.**
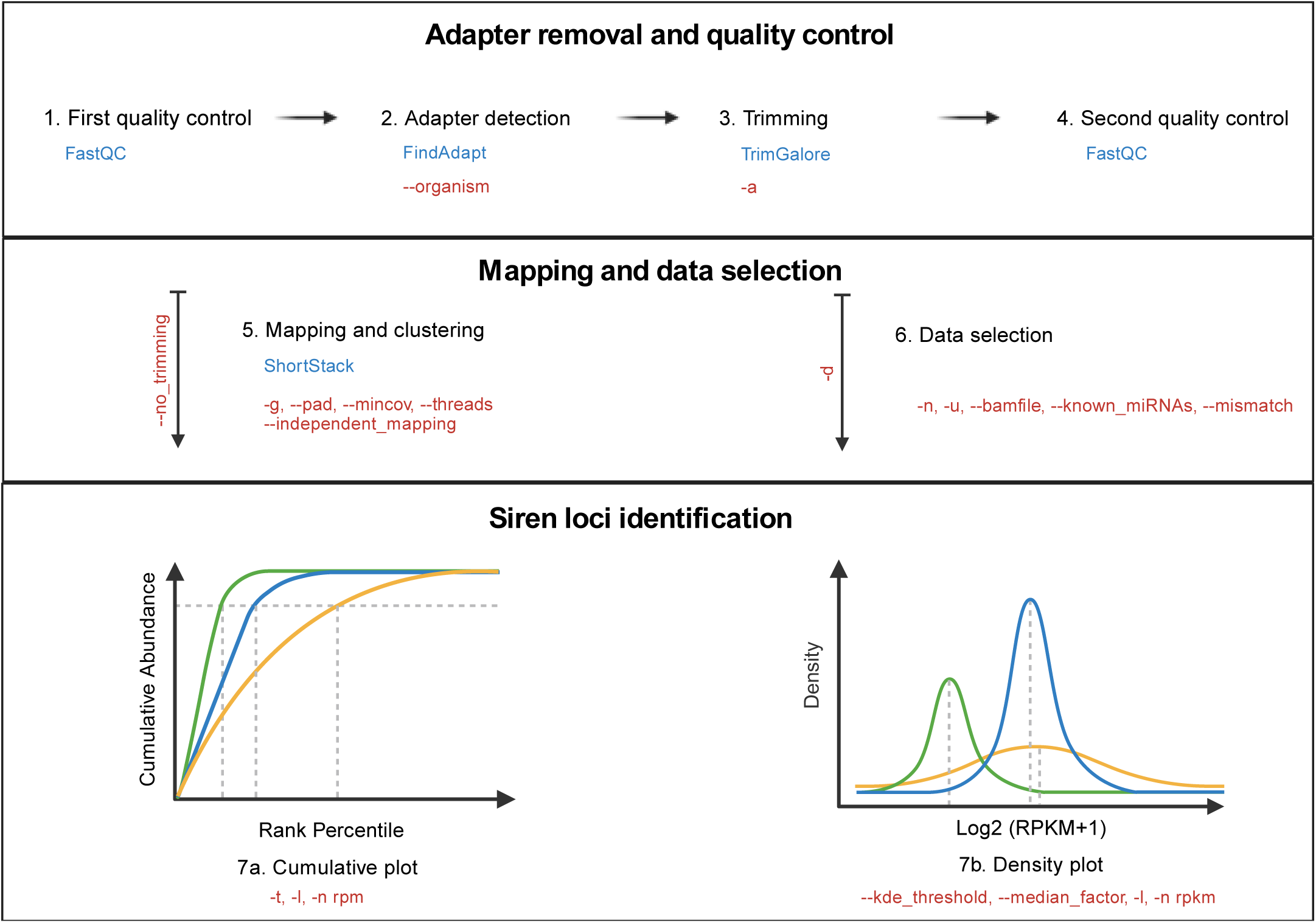
Workflow of SirenScan. The integrated bioinformatic tools are shown in blue and the arguments provided are shown in red. Sequencing quality control is done by FastQC (Andrews, 2010), the adapter detection by FindAdapt (Chen *et al*., 2024), the trimming by TrimGalore (Krueger, 2015) and Cutadapt (Martin, 2011) and the mapping and clustering by ShortStack (Axtell, 2013). For further technical aspects and other python libraries used by SirenScan, please refer to the <u>GitHub repository</u>.

### SirenScan enables robust identification of siren loci across datasets

To assess the reliability of SirenScan, we applied both cumulation-based and density-based strategies to four biological replicates of published *Arabidopsis* ovule sRNA sequencing data (Zhou *et al*., 2022). Using the cumulation-based approach with a 90% threshold, 142 siren loci were consistently identified across all replicates **(Supplementary Fig. 2a)**. In parallel, the density-based approach, based on local minimum detection, identified 128 loci shared across replicates **(Supplementary Fig. 2b, c)**. Notably, all density-based siren loci were contained within the cumulation-based set, whereas the additional 14 loci uniquely identified by the cumulation-based method were located at the boundary of the high-expression population in the density distribution **(Supplementary Fig. 2d, e)**. This indicates that the density-based approach is more stringent in defining highly expressed loci. Further characterization revealed that siren loci are not only among the most highly expressed sRNA–producing clusters but also significantly longer than other loci **(Supplementary Fig. 3a-d)**, a feature previously reported in *Brassica rapa* and potentially conserved across species (Grover *et al*., 2020). To further evaluate the consistency between different siren definitions and independent datasets, we compared siren loci identified by SirenScan (with both cumulative and density approaches) with three previously reported datasets, namely siren loci found in Arabidopsis ovules by Burgess *et al*., 2022 [18] and siren loci coupled with CLSY3 ChIP-seq peaks published in Zhou *et al*., 2022 (Zhou *et al*., 2022). Genomic intersections across datasets were visualized using UpSet plots **(Fig. 2a)**, revealing a substantial core of shared loci despite differences in identification strategies and coordinate definitions. Notably, 44% of genomic regions (73/165) were shared across all datasets, and this proportion increased to 76% when considering only the density-based, cumulation-based, and Zhou datasets, excluding smaller or less comparable datasets such as Burgess and CLSY3 ChIP-seq peaks. Length distributions across datasets were highly similar with the exception of Zhou’s data **(Fig. 2b)**, supporting the robustness of siren locus features. Genome-wide distributions showed consistent enrichment patterns across chromosomes **(Fig. 2c)**, indicating that the genomic regions identified as siren loci are reproducible. Quantitative comparison using Cobind (Ma *et al*., 2023) showed high overlap between datasets based on the Szymkiewicz-Simpson (SS) coefficient **(Fig. 2d)**. The cumulation-based and density-based approaches exhibited the highest similarity with 1.0 SS values, indicating strong agreement between the two strategies. Notably, high SS values were also observed in comparisons where one dataset is largely contained within another. This reflects the definition of the SS coefficient, which measures the proportion of overlap relative to the smaller dataset and therefore captures inclusion relationships between datasets. In contrast, Z-scores provide a composite measure of similarity that integrates multiple aspects of overlap. Together, these analyses demonstrate that while different siren identification strategies exhibit trade-offs between sensitivity and stringency, they converge on a largely shared set of high-confidence loci, supporting the robustness and reproducibility of SirenScan.

**Fig. 2.**
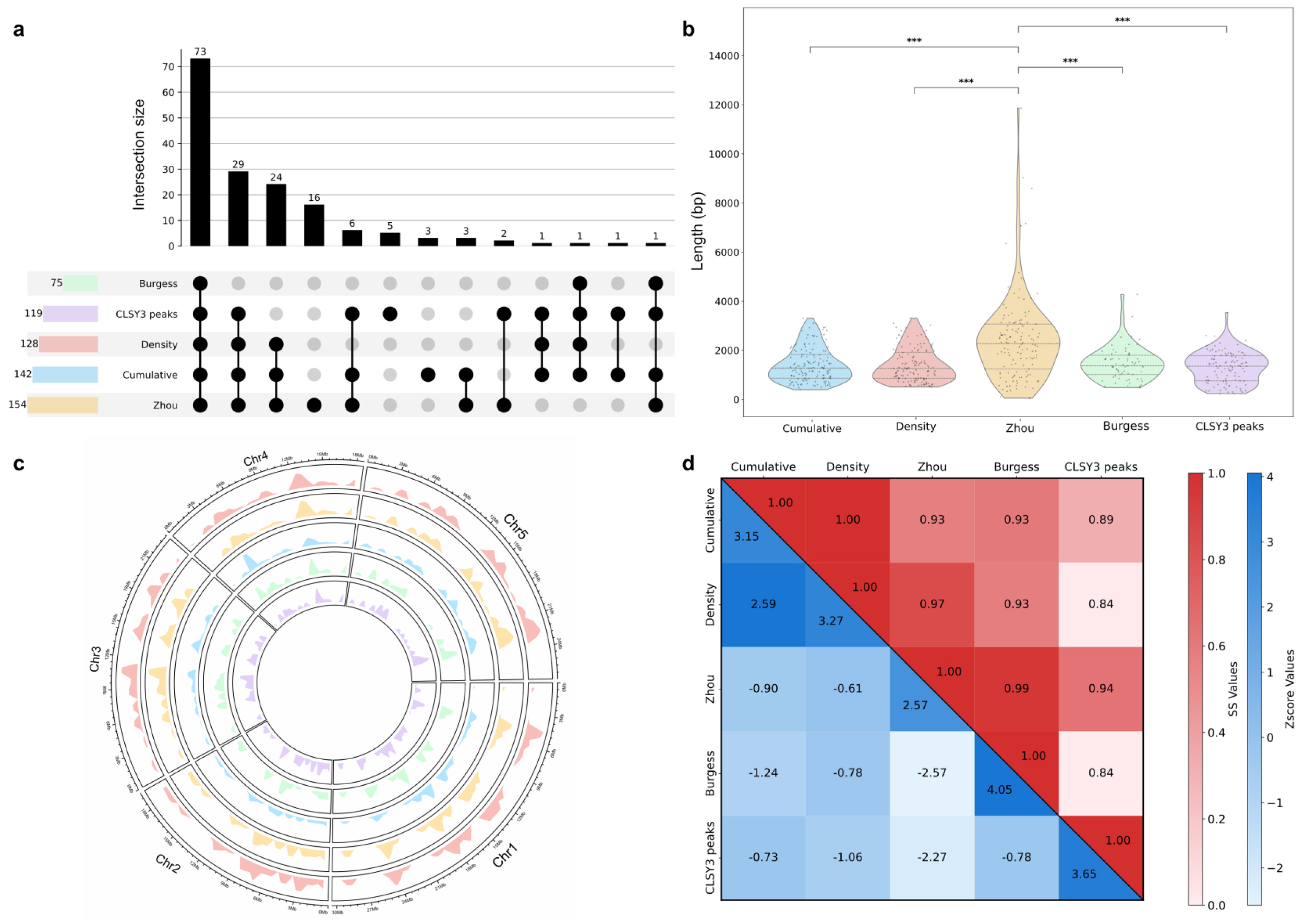
Validation of siren loci identified by SirenScan in *Arabidopsis thaliana* ovules. **a**. UpSet plot showing overlap of siren loci across datasets. The cumulative and density dataset are output by SirenScan. Burgess dataset from Burgess *et al*., 2022 (Burgess *et al*., 2022) and Zhou dataset with CLSY3 ChIP-seq peaks published in Zhou *et al*., 2022 (Zhou *et al*., 2022). Differences in dataset size reflect coordinate fragmentation or merging during cross-dataset comparisons. **b.** Length distribution of siren loci across datasets. Mann-Whitney U test with Bonferroni correction was applied (*** = p < 0.001). **c.** Genome-wide distribution of siren loci across the TAIR10 reference genome. Color code the same as in panel a. **d.** Pairwise comparison of genomic overlap using the Szymkiewicz–Simpson (SS) coefficient (in red) and associated Z-scores calculated (in blue) with Cobind (Ma *et al*., 2023).

### Siren RNAs are present across angiosperm ovules but exhibit lineage-specific expression dominance

Using SirenScan, we systematically screened for siren loci across sRNA sequencing datasets encompassing four additional plant species, namely *Capsella rubella*, *Oryza sativa* (rice), *Solanum lycopersicum* (tomato), and *Nymphaea colorata* (waterlily), including both publicly available and newly generated data **(Supplementary Table 1)**. Previous studies in *Arabidopsis* reported that siren RNAs are present in ovules and endosperm, but largely absent from vegetative tissues such as leaves (Rodrigues *et al*., 2013; Grover *et al*., 2020; Burgess *et al*., 2022; Zhou *et al*., 2022; Dziasek *et al*., 2024; Li *et al*., 2022; Rodrigues *et al*., 2021). To examine whether this pattern extends beyond *Arabidopsis*, we analyzed the other four species by comparing ovule tissues with vegetative tissues. SirenScan was applied to each dataset to detect siren loci and assess the presence and strength of siren RNA expression patterns across species. We first examined *Capsella rubella*, a close relative of *Arabidopsis thaliana*, which diverged roughly 10–14 million years ago (Koch & Kiefer, 2005). In ovule tissues, a small fraction of sRNA producing loci (∼5–6%) accounted for 90% of total sRNA abundance above 2 RPM, indicating a strong expression dominance pattern **(Fig. 3a)**. In contrast, more than 60% of loci were required to reach the same cumulative abundance in leaf tissues, reflecting a more uniform expression distribution. Consistently, density plots revealed a distinct population of highly expressed and longer loci exclusively in ovule samples, which was absent in leaves **(Fig. 3b)**. These observations indicate that the presence of siren loci in ovule tissues is conserved in *Capsella*. Building on previous studies showing the presence of siren RNAs in the endosperm of *Oryza sativa* ssp. *japonica* (Rodrigues *et al*., 2013), we next examined ovule tissues. Although approximately 40% of sRNA producing loci were required to account for 90% of total sRNA abundance, a much smaller fraction of loci (<1%) already contributed ∼70% of total abundance above 2 RPM in ovules. In contrast, ∼35% of loci were required to reach the same level in seedling tissues **(Fig. 3c)**. Consistently, density plots revealed a distinct population of highly expressed and longer loci exclusively in ovule samples, which was not observed in seedlings **(Fig. 3d)**. These results indicate that while siren-like expression patterns are present in rice ovules, the conventional 90% cumulative threshold may not be universally applicable across species. Instead, density-based approaches that capture distinct high-expression populations may provide a more robust measure for defining siren loci in such cases. In *Solanum*, the proportion of loci required to reach the 90% cumulative threshold was similar between ovule and leaf tissues, however, a clear difference in expression dominance was observed at lower thresholds. In ovules, only ∼2–3% of loci contributed to 40% of total sRNA abundance above 2 RPM, whereas more than 10% of loci were required to reach the same level in leaf tissues **(Fig. 3e)**. Consistently, density plots revealed a distinct population of highly expressed and longer loci exclusively in ovule samples, which was absent in leaves **(Fig. 3f)**. To further assess whether the siren locus phenomenon is a conserved feature in angiosperms, we examined sRNA profiles in the early-diverging angiosperm *Nymphaea colorata* (Tang *et al*., 2020). In ovule tissues, approximately 10–20% of loci accounted for 90% of total sRNA abundance above 2 RPM, whereas 60–70% of loci were required to reach the same threshold in leaf tissues **(Fig. 3g)**. Consistently, density plots revealed a distinct population of highly expressed and longer loci uniquely present in ovule samples, which was absent in leaves **(Fig. 3h)**. Taken together, these results demonstrate that the presence of siren loci in ovule tissues is a conserved feature across angiosperms, despite substantial variation in the strength and distribution of expression dominance among species. While the cumulative contribution of siren loci differs across evolutionary lineages, density-based analyses consistently reveal a distinct population of highly expressed loci in female reproductive tissues. This variability highlights the limitations of uniform thresholds and recommends the use of density-based approaches to robustly capture siren loci across diverse plant lineages, which treat each locus individually by intrinsic features rather than arbitrary requirements of aggregated patterns.

**Fig. 3.**
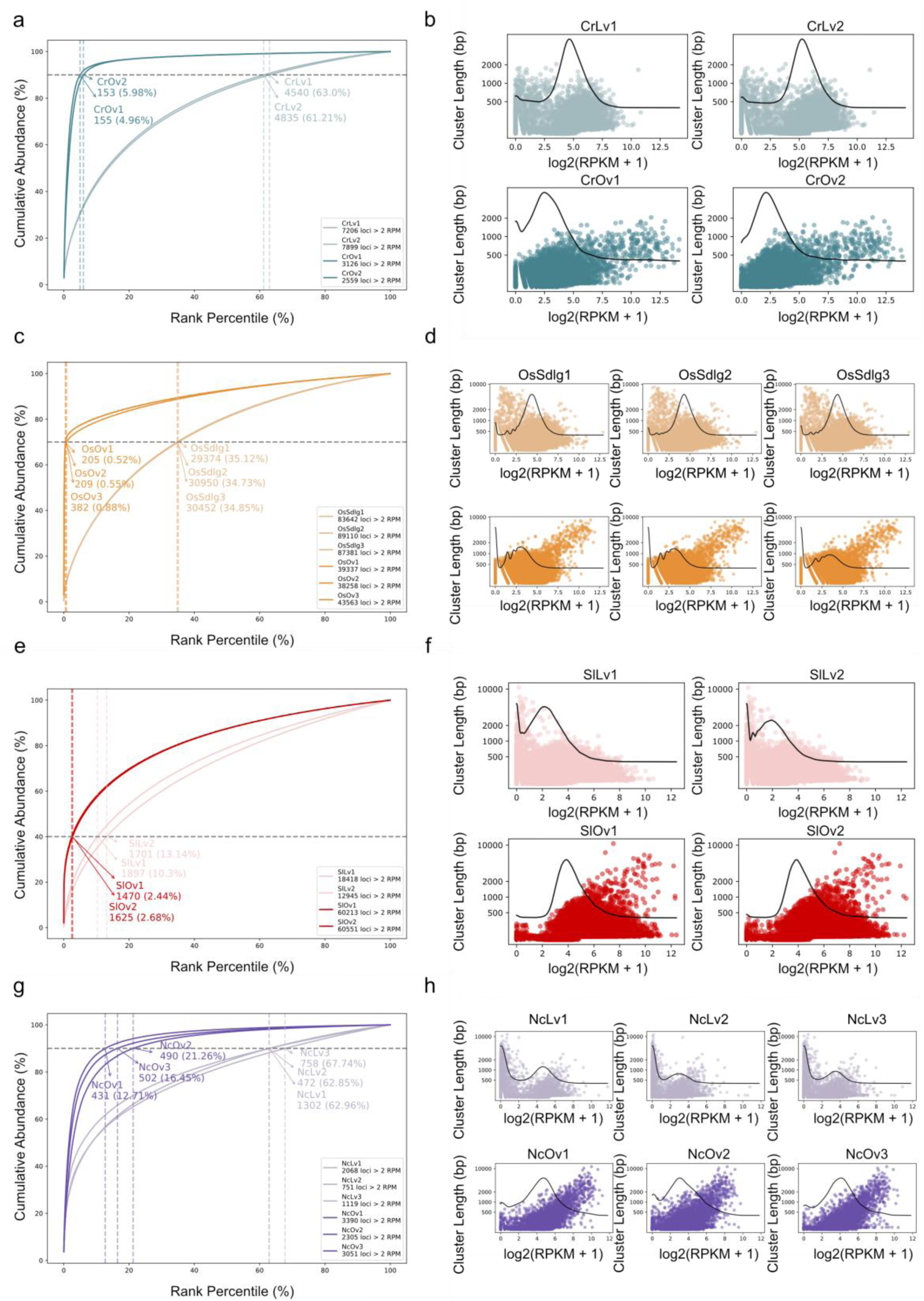
Identification of siren phenomenon across angiosperms using SirenScan. **a.–b**. Cumulation-based and density-based identification of siren loci in leaf and ovule tissues of *Capsella rubella*. **c.–d.** Cumulation-based and density-based identification in seedling and ovule tissues of *Oryza sativa*. **e.–f.** Cumulation-based and density-based identification in leaf and ovule tissues of *Solanum lycopersicum*. **g.–h.** Cumulation-based and density-based identification in leaf and ovule tissues of *Nymphaea colorata*.

### Lineage-specific diversification of TEs associated with siren loci

We analyzed ovule-specific siren loci in each species identified by the density-based approach implemented in SirenScan **(Supplementary Fig. 4)**. In *Capsella* and *Oryza*, expression density distributions exhibited clear bimodality, allowing siren loci to be defined using local minima as thresholds **(Supplementary Fig. 4a and 4c)**. The resulting loci showed near-omplete overlap with those identified using the cumulation-based approach **(Supplementary Fig. 4b and 4d)**, indicating strong concordance between methods. In *Solanum* and *Nymphaea*, density distributions lacked a clear bimodal structure **(Supplementary Fig. 4e and 4g)**, therefore, highly expressed loci were defined using a median-based threshold of log2(RPKM+1) values. To better capture the distinct high-expression populations in each species, thresholds were adjusted to 6× median for *Solanum* and *Nymphaea* including size filtering to capture the populations with high expression in ovules. Despite these differences, the resulting loci retained substantial overlap (> 94%) with those identified by the cumulation-based approach **(Supplementary Fig. 4f and 4h)**. Using this strategy, we identified 115, 104, 105, and 218 siren loci in *Capsella*, *Oryza*, *Solanum*, and *Nymphaea*, respectively.

**Fig. 4.**
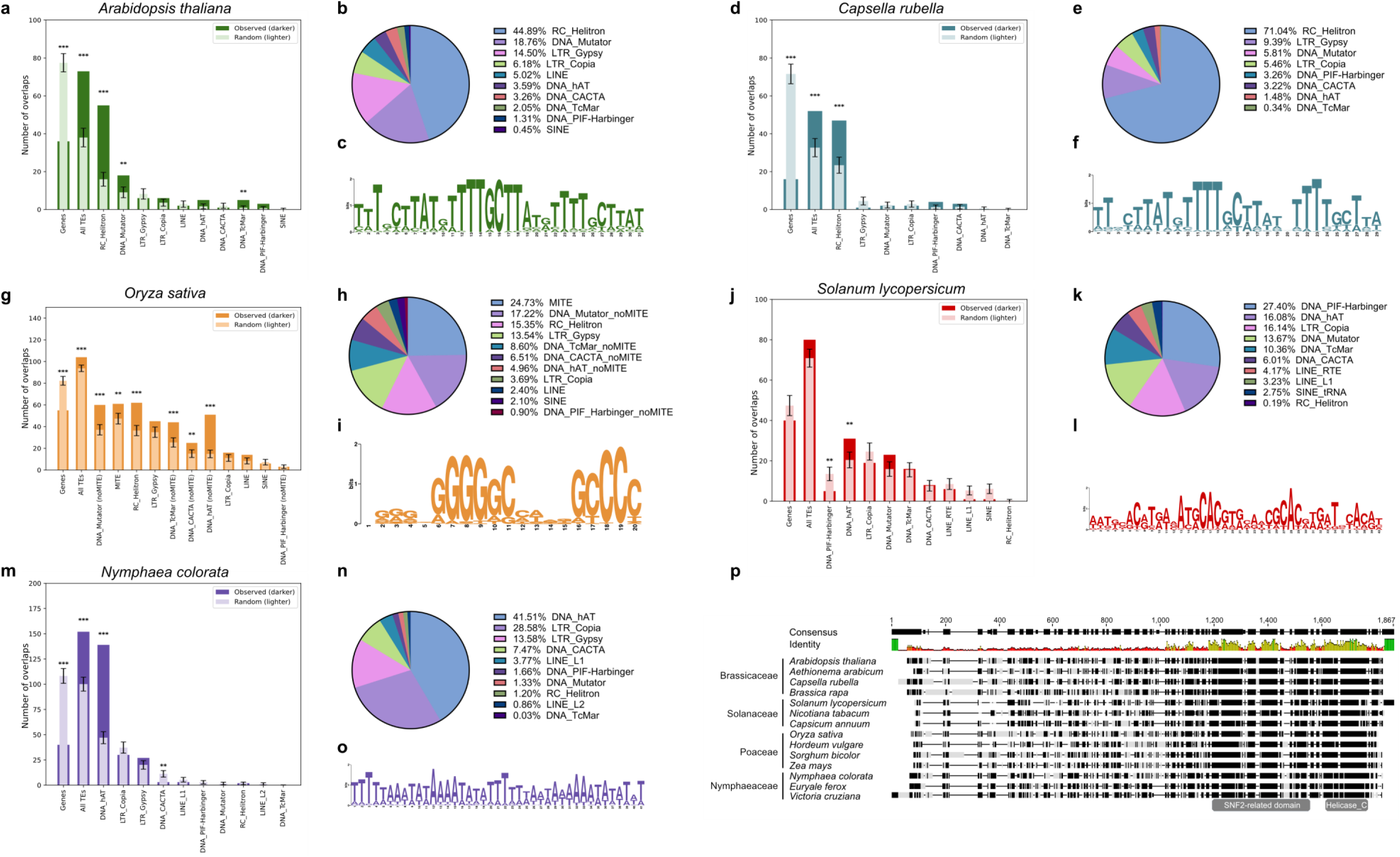
Regulatory and transposon-associated features of ovule siren loci across species. **a-c**. *Arabidopsis thaliana* **d-f.** *Capsella rubella* **g-i.** *Oryza sativa* **j-l.** *Solanum lycopersicum* **m-o.** *Nymphaea colorata*. **a,d,g,j,m.** Overlap between siren loci and genomic features. Dark bars represent observed overlaps between siren loci and TE superfamilies, whereas light bars represent randomized genomic regions used as controls. MITE elements were identified and separated from the DNA/TcMar, DNA/Mutator, DNA/CACTA, DNA/hAT, and DNA/PIF_Harbinger superfamilies, and the remaining elements within each superfamily were classified as non-MITEs. P-values were calculated using Monte Carlo permutation tests. (** = p < 0.01 and *** = p < 0.001 are shown). **b,e,h,k,n.** Relative abundance of TE superfamilies in each species genome. **c,f,i,l,o.** Motif enrichment analysis of siren loci in each species using MEME. **p.** Multiple sequence alignment of CLSY3/4 proteins from representative plant species, highlighting conservation of the SNF2 and Helicase-C domains and divergence within N-terminal regions. Residues shaded in black indicate positions with complete similarity based on the BLOSUM62 similarity matrix implemented in Geneious Prime.

To further investigate the genomic context of siren loci, we examined their overlap with genes and TE superfamilies. Across species, siren loci were generally enriched in TEs and depleted from genic regions **(Fig. 4a, 4d, 4g, 4j and 4m)**. However, the TE superfamilies associated with siren loci varied substantially across species. In *Arabidopsis* and *Capsella*, siren loci were strongly enriched in the RC/Helitron superfamily **(Fig. 4a and 4d)**. This is consistent with previous findings in *Brassica rapa*, where siren loci are associated with the Helitron *Persephone* family, which also contains AtCLSY3-like binding motifs (Burgess *et al*., 2022). In *Oryza*, siren loci were associated with diverse class II DNA transposons, including MITE-related families from the DNA/TcMar, DNA/Mutator, DNA/CACTA, and DNA/hAT superfamilies **(Fig. 4g)** (Soundiramourtty & Mirouze, 2025). In *Solanum*, sRNA production was largely restricted to chromosome arms **(Supplementary Fig. 5a)**, which correspond to euchromatic, gene-rich regions, in contrast to the pericentromeric enrichment observed in *Arabidopsis* **(Supplementary Fig.5b)**. This distribution is consistent with previous observations that in *Solanum* sRNA production is concentrated in chromosome arms rather than pericentromeric heterochromatin, correlating with spatial confinement of Pol IV activity (Sato *et al*., 2012; Corem *et al*., 2018). The most abundant TE superfamily in the tomato genome, LTR/Gypsy, is predominantly located in heterochromatic regions **(Fig. 4k and Supplementary Fig. 6a)** (Di Filippo *et al*., 2012) and significantly depleted from siren loci **(Supplementary Fig. 6b)**. Because the chromosomal distributions of siren loci and LTR/Gypsy element-containing pericentromeric regions are largely non-overlapping, we generated a euchromatin-restricted null model by excluding LTR/Gypsy regions from the permutation analysis **(Fig. 4j)**. This analysis revealed that siren loci showed significant enrichment for the DNA/hAT superfamily **(Fig. 4j)**. Similarly, in *Nymphaea*, siren loci were also primarily enriched in the DNA/hAT superfamily **(Fig. 4m)**. Together, these results indicate that siren loci are consistently associated with DNA TEs, although the specific TE superfamilies involved differ across lineages.

Interestingly, this variation reflects both TE abundance and genomic accessibility. In *Arabidopsis* and *Capsella*, Helitrons are the most abundant TEs by copy number **(Fig. 4b, 4e)** (Quesneville, 2020), suggesting a relationship between TE abundance and siren RNA production. Similarly, in *Oryza*, siren loci are enriched in TE superfamilies that give rise to non-autonomous MITEs **(Fig. 4g)** (Soundiramourtty & Mirouze, 2025), which are among the most abundant TEs in rice **(Fig. 4h)** (Turcotte *et al*., 2001). In *Nymphaea*, siren loci are associated with DNA/hAT elements **(Fig. 4m)**, the most abundant DNA transposons in this genome **(Fig. 4n)** (Zhang *et al*., 2020b). By contrast, in *Solanum*, LTR/Gypsy elements, despite representing the most abundant TE superfamily in the genome, were not enriched, likely because they are predominantly localized in heterochromatic regions. Instead, siren loci are enriched in TE superfamilies that are highly abundant on chromosome arms (Su *et al*., 2021), consistent with the greater accessibility of these regions to Pol IV (Corem *et al*., 2018). Thus, siren RNAs appear to be preferentially generated from TE superfamilies that are either highly abundant or accessible to the Pol IV machinery within each genome **(Fig. 4k and Supplementary Fig. 6a, c)**, reflecting a conserved association with TEs alongside lineage-specific diversification.

### Siren loci are enriched for lineage-specific sequence motifs

Previous studies in *Arabidopsis* have shown that siren loci are enriched for sequence motifs corresponding to AtCLSY3 binding sites (Zhou *et al*., 2022; Wu *et al*., 2025; Xu *et al*., 2025). To investigate whether such regulatory features are conserved or diversified across species, we next performed a motif enrichment analysis of siren loci using MEME (Bailey *et al*., 2015). The top-ranked motifs are shown in **(Fig. 4c, 4f, 4i, 4l and 4o)** and other motifs are shown in **(Supplementary Fig. 7)**. In *Arabidopsis* and *Capsella*, siren loci were enriched for highly similar motifs that match the known AtCLSY3 binding site (Zhou *et al*., 2022; Wu *et al*., 2025; Xu *et al*., 2025), indicating conservation of *cis*-regulatory features within *Brassicaceae*. In contrast, other species exhibited more divergent motif patterns, suggesting lineage-specific regulatory diversification of siren loci.

Given that AtCLSY3 and AtCLSY4 function in association with REM transcription factors and are essential for recruiting Pol IV to siren loci (Wu *et al*., 2025; Xu *et al*., 2025; Pandesha & Slotkin, 2025), we hypothesized that divergence of CLSY3/4 proteins across species may contribute to the observed variation in motif enrichment. To explore this possibility, we performed sequence alignment of CLSY3/4 orthologs across species **(Fig. 4p and Supplementary Fig. 8)**. We found that the SNF2 and Helicase-C domains are highly conserved, consistent with their core roles in chromatin remodeling and Pol IV recruitment. In contrast, the N-terminal regions show substantial sequence divergence among species from different families. This pattern suggests that the N-terminal region may mediate lineage-specific interactions, potentially with distinct TFs recognizing different motifs, thereby contributing to lineage-specific targeting of Pol IV and the diversification of siren locus regulation.

## Discussion

From a methodological perspective, our results highlight the limitations of defining siren loci using fixed cumulative thresholds. While cumulation-based approaches capture the overall dominance of sRNA production, they may fail to identify distinct high-expression populations in species with weaker dominance patterns. By integrating rank–abundance and density-based analyses, SirenScan provides a more flexible and data-driven pipeline that adapts to species-specific expression landscapes. This approach enables robust identification and comparison of siren loci across diverse biological contexts.

Siren RNAs were originally described as a small subset of loci producing a disproportionate fraction of sRNAs in reproductive tissues (Rodrigues *et al*., 2013). While this phenomenon has been well characterized in a few model systems (Grover *et al*., 2020; Zhou *et al*., 2022; Burgess *et al*., 2022; Dziasek *et al*., 2024), its broader evolutionary distribution and regulatory basis have remained unclear. By applying SirenScan across angiosperm phylogeny, we show that the presence of siren loci in ovule tissues is a conserved feature of flowering plants, even extending to early diverging angiosperms **(Fig. 3)**. These findings suggest that siren RNA production represents a fundamental aspect of reproductive sRNA landscapes in angiosperms. Despite this conserved presence, we observed substantial variation in the strength and distribution of expression dominance across species. In some lineages, such as *Brassicaceae* and *Nymphaea*, a small fraction of loci accounts for the vast majority of sRNA abundance **(Fig.3 a-b, g-h)**, whereas in others, including *Oryza* and *Solanum*, this dominance is more attenuated **(Fig.3 c-f)**. This variability likely reflects differences in genome organization and chromatin accessibility across species.

Our analyses further reveal that siren loci are consistently associated with TEs, although the specific TE families involved vary across lineages. In *Brassicaceae*, siren loci are strongly enriched in Helitron elements, consistent with previous observations (Burgess *et al*., 2022), whereas in other species they are associated with a broader range of DNA transposon superfamilies. Nevertheless, what they share in common is that TEs associated with siren loci are highly abundant DNA TE families, suggesting that siren RNA formation is connected to the silencing of DNA TEs. At the regulatory level, motif analysis indicates that siren loci in *Brassicaceae* share conserved *cis*-regulatory features resembling CLSY3 binding sites, whereas other species exhibit other lineage-specific motifs divergent from each other. Together with the observed divergence in the N-terminal region of CLSY3/4 protein sequences, these results suggest that siren locus specification is mediated by a conserved core machinery combined with lineage-specific targeting mechanisms. We propose that conserved C-terminal domains of CLSY3/4 proteins maintain core Pol IV recruitment functions, while N-terminal variable regions enable interactions with distinct TFs, thereby shaping species-specific siren locus targeting. Consistent with this model, previous studies have shown that OsCLSY3 and OsCLSY4 are unable to fully complement Arabidopsis *clsy* mutants in restoring DNA methylation patterns, supporting functional divergence of CLSY3/4 proteins across species (Xu *et al*., 2024). Why different species may employ different TFs for CLSY recruitment and TE silencing remains an exciting question of future research. Perhaps, species-specific expansion of distinct TE families drove the expansion of specific TFs recognizing these TEs, similar to the species-specific recognition of TEs by zinc-finger protein (ZFP) TFs in mammals (Jacobs *et al*., 2014; Yang *et al*., 2017). Rapidly evolving KRAB-ZFPs bind lineage-specific TE motifs and recruit repressive chromatin machinery, similar to the recruitment of Pol IV by CLSY3 and REM transcription factors in *Arabidopsis*.

In summary, our study establishes siren RNA expression as a conserved yet dynamically evolving feature of angiosperm female reproductive tissues and generates novel insights into the evolution of sRNA landscapes in angiosperms.

## Data availability

Small RNA sequencing data generated in this study are available in the NCBI under the project PRJNA1465687. SirenScan is available under the MIT license on GitHub: https://github.com/AurelienVALENTIN/SirenScan/. Data to prepare figures are available at https://zenodo.org/records/21165642.

## Acknowledgements

We thank Botanischen Garten der Universität Potsdam and D. Barro-Trastoy for providing the *Nymphaea colorata* materials.

## Contributions

H.P., A.V., Y.Q. and C.K. conceptualized the project, developed the methodology and provided supervision. H.P., A.V. and K.D. conducted experiments. H.P., A.V. and Y.Q. performed bioinformatic analyses. C.K. acquired funding and administered the project. H.P., A.V. and C.K. wrote the original paper draft. All authors reviewed and edited the paper.

## Reference

1. Andrews S. 2010. FastQC: A Quality Control Tool for High Throughput Sequence Data [Online].

2. Axtell MJ. 2013. ShortStack: Comprehensive annotation and quantification of small RNA genes. RNA 19: 740–751.

3. Bailey TL, Johnson J, Grant CE, Noble WS. 2015. The MEME suite. Nucleic Acids Research 43: W39– W49.

4. Batista RA, Moreno-Romero J, Qiu Y, van Boven J, Santos-González J, Figueiredo DD, Köhler C. 2019. The MADS-box transcription factor PHERES1 controls imprinting in the endosperm by binding to domesticated transposons (D Zilberman and CS Hardtke, Eds). eLife 8: e50541.

5. Bélanger S, Zhan J, Meyers BC. 2023. Phylogenetic analyses of seven protein families refine the evolution of small RNA pathways in green plants. Plant Physiology 192: 1183–1203.

6. Burgess D, Chow HT, Grover JW, Freeling M, Mosher RA. 2022. Ovule siRNAs methylate protein-coding genes in trans. The Plant Cell 34: 3647–3664.

7. Chakraborty T, Trujillo JT, Kendall T, Mosher RA. 2024. Charophytic green algae encode ancestral polymerase IV/polymerase V subunits and a CLSY/DRD1 homolog. Genome Biology and Evolution 16: evae119.

8. Chen Y, Tian J, Zhao Y, Zhang J, Liang C. 2026. A telomere-to-telomere reference genome assembly of tomato cultivar ‘Heinz 1706’. Plant Communications 7: 101618.

9. Chen H-C, Wang J, Shyr Y, Liu Q. 2024. FindAdapt: A python package for fast and accurate adapter detection in small RNA sequencing. PLOS Computational Biology 20: e1011786.

10. Corem S, Doron-Faigenboim A, Jouffroy O, Maumus F, Arazi T, Bouché N. 2018. Redistribution of CHH methylation and small interfering RNAs across the genome of tomato ddm1 mutants. The Plant Cell 30: 1628–1644.

11. Cuerda-Gil D, Slotkin RK. 2016. Non-canonical RNA-directed DNA methylation. Nature Plants 2.

12. Di Filippo M, Traini A, D’Agostino N, Frusciante L, Chiusano ML. 2012. Euchromatic and heterochromatic compositional properties emerging from the analysis of Solanum lycopersicum BAC sequences. Gene 499: 176–181.

13. Dziasek K, Santos-González J, Wang K, Qiu Y, Zhu J, Rigola D, Nijbroek K, Köhler C. 2024. Dosage-sensitive maternal siRNAs determine hybridization success in Capsella. Nature Plants 10: 1969–1983.

14. Edgar RC. 2004. MUSCLE: multiple sequence alignment with high accuracy and high throughput. Nucleic Acids Research 32: 1792–1797.

15. Erdmann RM, Satyaki PRV, Klosinska M, Gehring M. 2017. A small RNA pathway mediates allelic dosage in endosperm. Cell Reports 21: 3364–3372.

16. Felgines L, Rymen B, Martins LM, Xu G, Matteoli C, Himber C, Zhou M, Eis J, Coruh C, Böhrer M, et al. 2024. CLSY docking to Pol IV requires a conserved domain critical for small RNA biogenesis and transposon silencing. Nature Communications 15: 10298.

17. Gel B, Díez-Villanueva A, Serra E, Buschbeck M, Peinado MA, Malinverni R. 2016. regioneR: an R/Bioconductor package for the association analysis of genomic regions based on permutation tests. Bioinformatics 32: 289–291.

18. Grover JW, Burgess D, Kendall T, Baten A, Pokhrel S, King GJ, Meyers BC, Freeling M, Mosher RA. 2020. Abundant expression of maternal siRNAs is a conserved feature of seed development. Proceedings of the National Academy of Sciences 117: 15305–15315.

19. Grover JW, Kendall T, Baten A, Burgess D, Freeling M, King GJ, Mosher RA. 2018. Maternal components of RNA-directed DNA methylation are required for seed development in Brassica rapa. The Plant Journal 94: 575–582.

20. Herr AJ, Jensen MB, Dalmay T, Baulcombe DC. 2005. RNA Polymerase IV Directs Silencing of Endogenous DNA. Science 308: 118–120.

21. Jacobs FMJ, Greenberg D, Nguyen N, Haeussler M, Ewing AD, Katzman S, Paten B, Salama SR, Haussler D. 2014. An evolutionary arms race between KRAB zinc-finger genes ZNF91/93 and SVA/L1 retrotransposons. Nature 516: 242–245.

22. Kim T, Resende MFR, Zhao M, Begcy K. 2024. A female-specific RdDM-associated element guides de novo DNA methylation during rice gametophyte development. bioRxiv.

23. Kirkbride RC, Lu J, Zhang C, Mosher RA, Baulcombe DC, Chen ZJ. 2019. Maternal small RNAs mediate spatial-temporal regulation of gene expression, imprinting, and seed development in Arabidopsis. Proceedings of the National Academy of Sciences 116: 2761–2766.

24. Koch MA, Kiefer M. 2005. Genome evolution among cruciferous plants: a lecture from the comparison of the genetic maps of three diploid species—Capsella rubella, Arabidopsis lyrata subsp. petraea, and A. thaliana. American Journal of Botany 92: 761–767.

25. Krueger F. 2015. Trim Galore!: A wrapper around Cutadapt and FastQC to consistently apply adapter and quality trimming to FastQ files, with extra functionality for RRBS data.

26. Lamesch P, Berardini TZ, Li D, Swarbreck D, Wilks C, Sasidharan R, Muller R, Dreher K, Alexander DL, Garcia-Hernandez M, et al. 2012. The Arabidopsis Information Resource (TAIR): improved gene annotation and new tools. Nucleic Acids Research 40: D1202–D1210.

27. Langmead B, Trapnell C, Pop M, Salzberg SL. 2009. Ultrafast and memory-efficient alignment of short DNA sequences to the human genome. Genome Biology 10: R25.

28. Li C, Gent JI, Xu H, Fu H, Russell SD, Sundaresan V. 2022. Resetting of the 24-nt siRNA landscape in rice zygotes. Genome Research 32: 309–323.

29. Lu J, Zhang C, Baulcombe DC, Chen ZJ. 2012. Maternal siRNAs as regulators of parental genome imbalance and gene expression in endosperm of *Arabidopsis* seeds. Proceedings of the National Academy of Sciences 109: 5529–5534.

30. Ma T, Guo L, Yan H, Wang L. 2023. Cobind: quantitative analysis of the genomic overlaps. Bioinformatics Advances 3: vbad104.

31. Martin M. 2011. Cutadapt removes adapter sequences from high-throughput sequencing reads. EMBnet.journal 17: 10–12.

32. Matzke MA, Mosher RA. 2014. RNA-directed DNA methylation: An epigenetic pathway of increasing complexity. Nature Reviews Genetics 15: 394–408.

33. Mosher RA, Melnyk CW, Kelly KA, Dunn RM, Studholme DJ, Baulcombe DC. 2009. Uniparental expression of PolIV-dependent siRNAs in developing endosperm of Arabidopsis. Nature 460: 283–286.

34. Pal AK, Gandhivel VH-S, Nambiar AB, Shivaprasad PV. 2024. Upstream regulator of genomic imprinting in rice endosperm is a small RNA-associated chromatin remodeler. Nature Communications 15: 7807.

35. Pal AK, Rana S, Dey R, Shivaprasad PV. 2025. Loss of function of chromatin remodeler OsCLSY4 leads to RdDM-mediated mis-expression of endosperm-specific genes affecting grain qualities. PLOS Genetics 21: e1011956.

36. Pandesha P, Slotkin RK. 2025. Transcription factor-mediated recruitment of small interfering RNA production. Nature Plants 11: 2453–2454.

37. Petrella R, Cucinotta M, Mendes MA, Underwood CJ, Colombo L. 2021. The emerging role of small RNAs in ovule development, a kind of magic. Plant Reproduction 34: 335–351.

38. Quesneville H. 2020. Twenty years of transposable element analysis in the Arabidopsis thaliana genome. Mobile DNA 11: 1–13.

39. Rodrigues JA, Hsieh P-H, Ruan D, Nishimura T, Sharma MK, Sharma R, Ye X, Nguyen ND, Nijjar S, Ronald PC, et al. 2021. Divergence among rice cultivars reveals roles for transposition and epimutation in ongoing evolution of genomic imprinting. Proceedings of the National Academy of Sciences 118: e2104445118.

40. Rodrigues JA, Ruan R, Nishimura T, Sharma MK, Sharma R, Ronald PC, Fischer RL, Zilberman D. 2013. Imprinted expression of genes and small RNA is associated with localized hypomethylation of the maternal genome in rice endosperm. Proceedings of the National Academy of Sciences 110: 7934– 7939.

41. Sato S, Tabata S, Hirakawa H, Asamizu E, Shirasawa K, Isobe S, Kaneko T, Nakamura Y, Shibata D, Aoki K, et al. 2012. The tomato genome sequence provides insights into fleshy fruit evolution. Nature 485: 635–641.

42. Shang L, He W, Wang T, Yang Y, Xu Q, Zhao X, Yang L, Zhang H, Li X, Lv Y, et al. 2023. A complete assembly of the rice Nipponbare reference genome. Molecular Plant 16: 1232–1236.

43. Slotte T, Hazzouri KM, Ågren JA, Koenig D, Maumus F, Guo Y-L, Steige K, Platts AE, Escobar JS, Newman LK, et al. 2013. The Capsella rubella genome and the genomic consequences of rapid mating system evolution. Nature Genetics 45: 831–835.

44. Soundiramourtty A, Mirouze M. 2025. Plant MITEs: miniature transposable elements with major impacts. Mobile DNA 16: 43.

45. Su X, Wang B, Geng X, Du Y, Yang Q, Liang B, Meng G, Gao Q, Yang W, Zhu Y, et al. 2021. A high-continuity and annotated tomato reference genome. BMC Genomics 22: 898.

46. Tang H, Zhang L, Chen F, Zhang X, Chen F, Ma H, Van de Peer Y. 2020. Nymphaea colorata (Blue-Petal Water Lily). Trends in Genetics 36: 718–719.

47. Trujillo JT, Seetharam AS, Hufford MB, Beilstein MA, Mosher RA. 2018. Evidence for a unique DNA-dependent RNA polymerase in cereal crops. Molecular Biology and Evolution 35: 2454–2462.

48. Turcotte K, Srinivasan S, Bureau T. 2001. Survey of transposable elements from rice genomic sequences. The Plant Journal 25: 169–179.

49. Vaucheret H, Voinnet O. 2024. The plant siRNA landscape. The Plant Cell 36: 246–275.

50. Wierzbicki AT, Haag JR, Pikaard CS. 2008. Noncoding transcription by RNA polymerase Pol IVb/Pol V mediates transcriptional silencing of overlapping and adjacent genes. Cell 135: 635–648.

51. Wu Z, Xue Y, Wang S, Shih Y-H, Zhong Z, Feng S, Draper J, Lu A, Hoeke CA, Sha J, et al. 2025. REM transcription factors and GDE1 shape the DNA methylation landscape through the recruitment of RNA polymerase IV transcription complexes. Nature Cell Biology 27: 1136–1147.

52. Xie Z, Johansen LK, Gustafson AM, Kasschau KD, Lellis AD, Zilberman D, Jacobsen SE, Carrington JC. 2004. Genetic and functional diversification of small RNA pathways in plants. PLOS Biology 2: e104.

53. Xu G, Chen Y, Martins LM, Li E, Wang F, Magana T, Ruan J, Law JA. 2025. Transcription factors instruct DNA methylation patterns in plant reproductive tissues. Nature Cell Biology.

54. Xu D, Zeng L, Wang L, Yang D-L. 2024. Rice requires a chromatin remodeler for Polymerase IV-small interfering RNA production and genomic immunity. Plant Physiology 194: 2149–2164.

55. Yang P, Wang Y, Macfarlan TS. 2017. The role of KRAB-ZFPs in transposable element repression and mammalian evolution. Transposable Elements 33: 871–881.

56. Zhang R, Chen Y, Xu M, Zhang Y, Liu Y, Wang L, Liu J-X, Hong L, Yang Y, Zhou M. 2026. OsCLSY4 modulates epigenomic patterns and grain size in rice. The Plant Journal 125: e70756.

57. Zhang L, Chen F, Zhang X, Li Z, Zhao Y, Lohaus R, Chang X, Dong W, Ho SYW, Liu X, et al. 2020a. The water lily genome and the early evolution of flowering plants. Nature 577: 79–84.

58. Zhang L, Chen F, Zhang X, Li Z, Zhao Y, Lohaus R, Chang X, Dong W, Ho SYW, Liu X, et al. 2020b. The water lily genome and the early evolution of flowering plants. Nature 577: 79–84.

59. Zhou M, Coruh C, Xu G, Martins LM, Bourbousse C, Lambolez A, Law JA. 2022. The CLASSY family controls tissue-specific DNA methylation patterns in Arabidopsis. Nature Communications 13: 244.

60. Zhou M, Palanca AMS, Law JA. 2018. Locus-specific control of the de novo DNA methylation pathway in Arabidopsis by the CLASSY family. Nature Genetics 50: 865–873.

61. Zilberman D, Cao X, Jacobsen SE. 2003. ARGONAUTE4 control of locus-specific siRNA accumulation and DNA and histone methylation. Science 299: 716–719.

