## Supplemental Figures for "SirenScan reveals the conserved presence of siren RNAs and their evolutionary diversification across angiosperms"

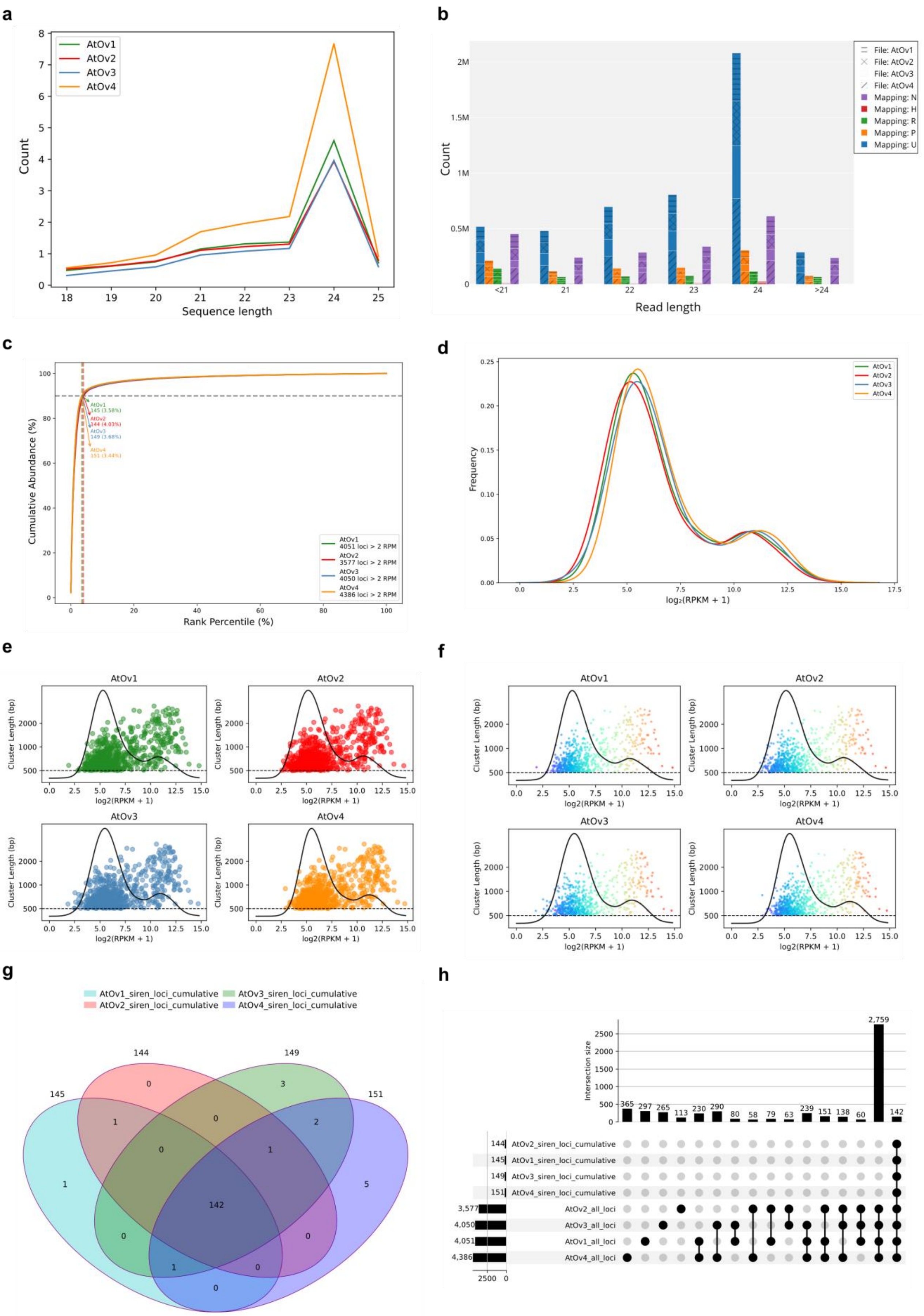
a
AtOv1
AtOv2
AtOv3
AtOv4
Count
Sequence length
b
File: AtOv1
File: AtOv2
File: AtOv3
File: AtOv4
Mapping: N
Mapping: H
Mapping: R
Mapping: P
Mapping: U
Count
Read length
c
Cumulative Abundance (%)
Rank Percentile (%)
AtOv1 4051 loci > 2 RPM
AtOv2 3577 loci > 2 RPM
AtOv3 4050 loci > 2 RPM
AtOv4 4386 loci > 2 RPM
d
Frequency
log₂(RPKM + 1)
e
f
Cluster Length (bp)
log2(RPKM + 1)
g
AtOv1_siren_loci_cumulative
AtOv2_siren_loci_cumulative
AtOv3_siren_loci_cumulative
AtOv4_siren_loci_cumulative
h
Intersection size
AtOv2_siren_loci_cumulative
AtOv1_siren_loci_cumulative
AtOv3_siren_loci_cumulative
AtOv4_siren_loci_cumulative
AtOv2_all_loci
AtOv3_all_loci
AtOv1_all_loci
AtOv4_all_loci

**Supplemental Fig.1 Representative outputs generated by SirenScan.** Representative plots generated by SirenScan using *Arabidopsis* ovule small RNA sequencing data (Zhou *et al.*, 2022). **a.** Small RNA length distribution after trimming per million reads. **b.** Read mapping categories assigned by ShortStack (Axtell, 2013). **c.** Cumulation-based (rank–abundance) identification of siren loci. **d.** Density-based identification of highly expressed loci (clusters <500 bp excluded). **e.** Scatter plot of cluster expression from panel d. **f.** Color-consistent scatter plot enabling tracking of individual clusters across samples. **g.** Venn diagram summarizing shared siren loci between samples. **h.** UpSet plot showing overlap of all loci across samples (intersections <1% omitted).

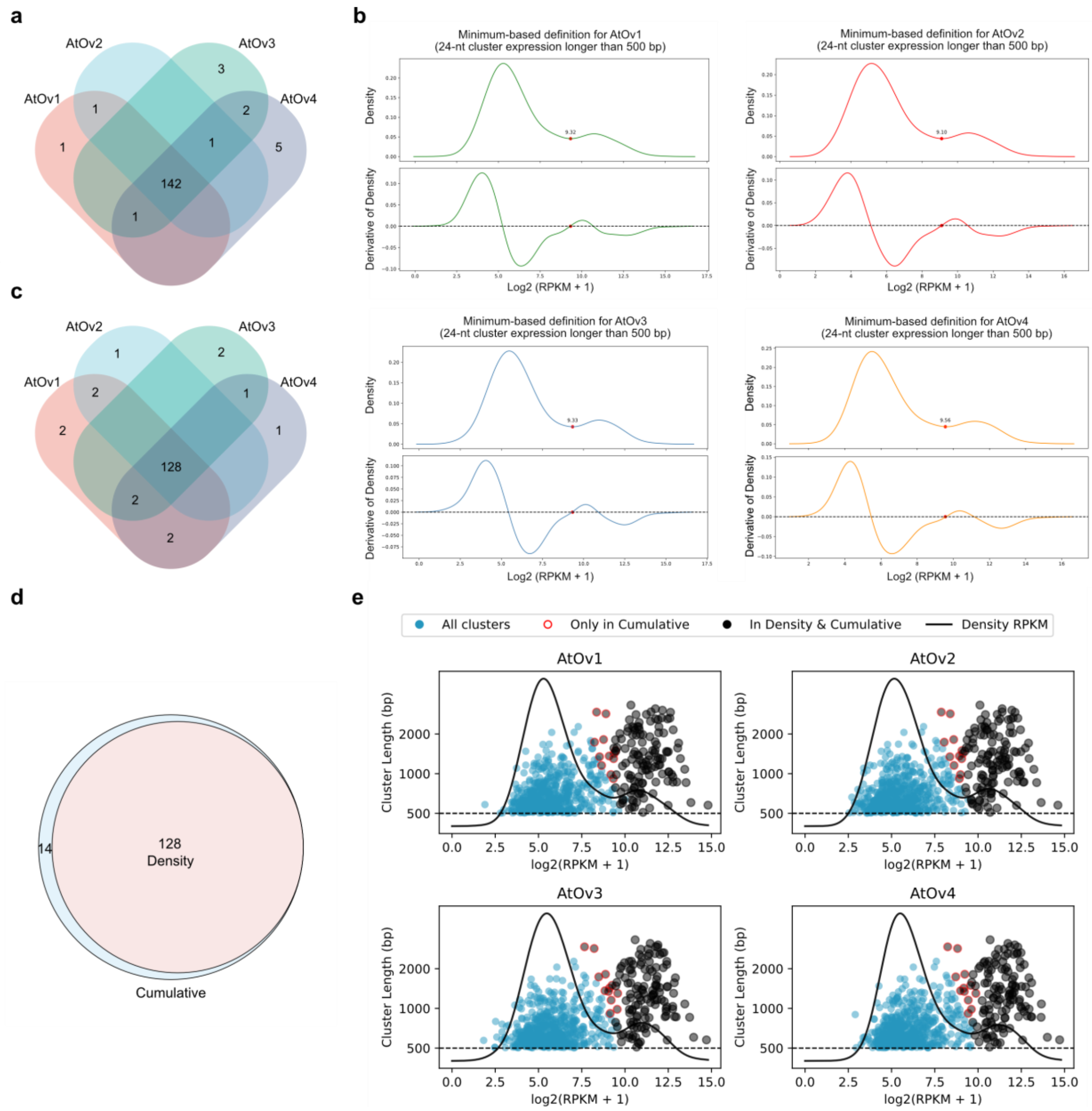


**Supplemental Fig.2 Comparison of cumulation- and density-based identification of siren loci a.** Overlap of siren loci identified across four biological replicates from Zhou *et al.*, 2022 (Zhou *et al.*, 2022) using the cumulation-based approach (90% threshold). **b.** Density plots and corresponding derivative curves showing the local minima used to define siren loci in each replicate. **c.** Overlap of siren loci identified across four biological replicates using the density-based approach. **d.** Overlap between siren loci identified by the cumulation-based and density-based approaches. **e.** Mapping of siren loci identified by both approaches onto the expression density distribution.

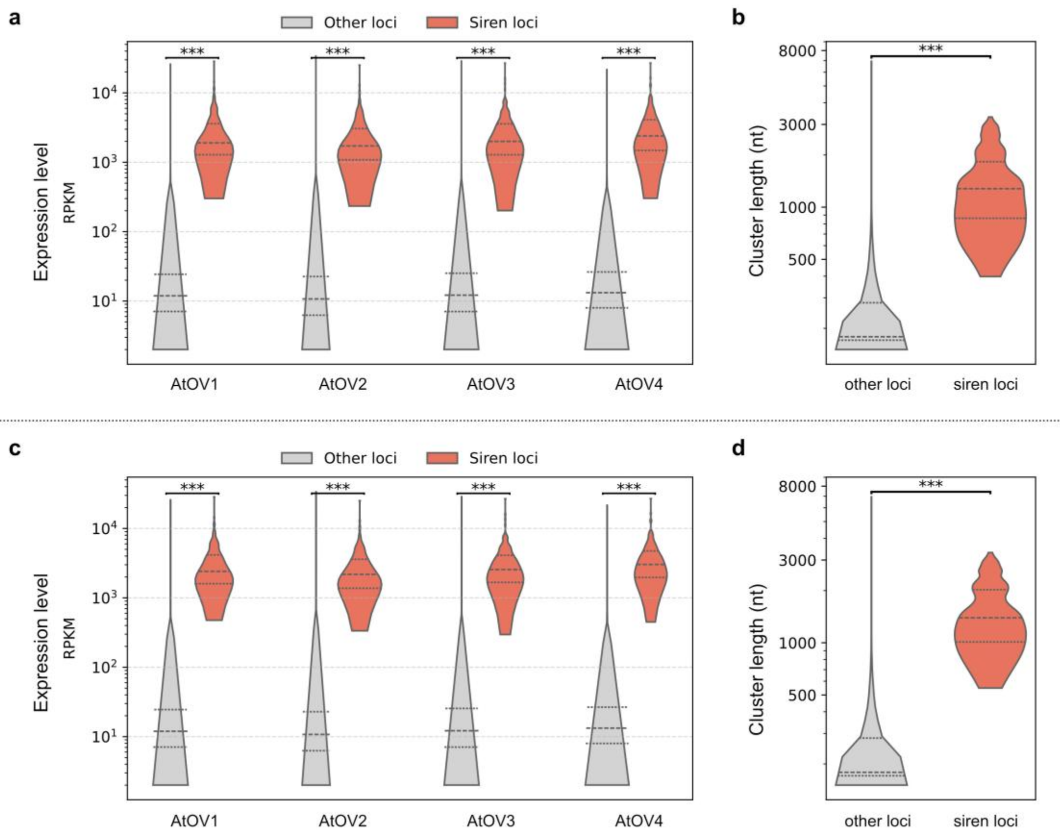


**Supplemental Fig.3 Distinct expression and length characteristics of siren loci a.** Expression levels of siren and non-siren loci using the cumulation-based approach. **b.** Cluster length of 142 common siren and non-siren loci using the cumulation-based approach. **c.** Expression levels of siren and non-siren loci using the density-based approach. **d.** Cluster length of 128 common siren and non-siren loci using the density-based approach. Mann–Whitney U test with Bonferroni correction was applied.

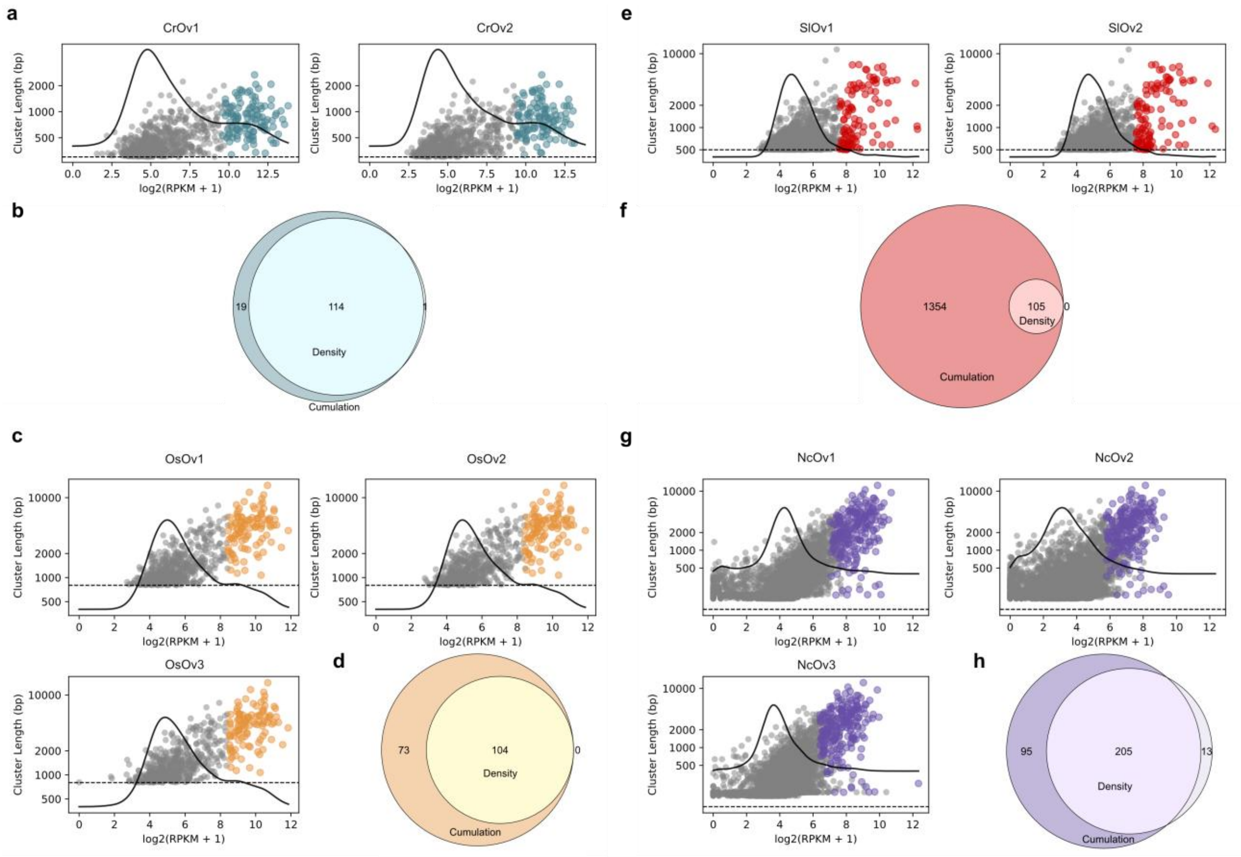


**Supplementary Fig.4. Identification of ovule siren loci across species using SirenScan. a,c,e,g.** Density-based identification of ovule siren loci in *Capsella rubella*, *Oryza sativa*, *Solanum lycopersicum*, and *Nymphaea colorata*. Grey dots represent non-siren loci, whereas colored dots indicate loci classified as siren loci. **b,d,f,h.** Venn diagrams showing the overlap between siren loci identified using cumulation-based and density-based approaches in each species.

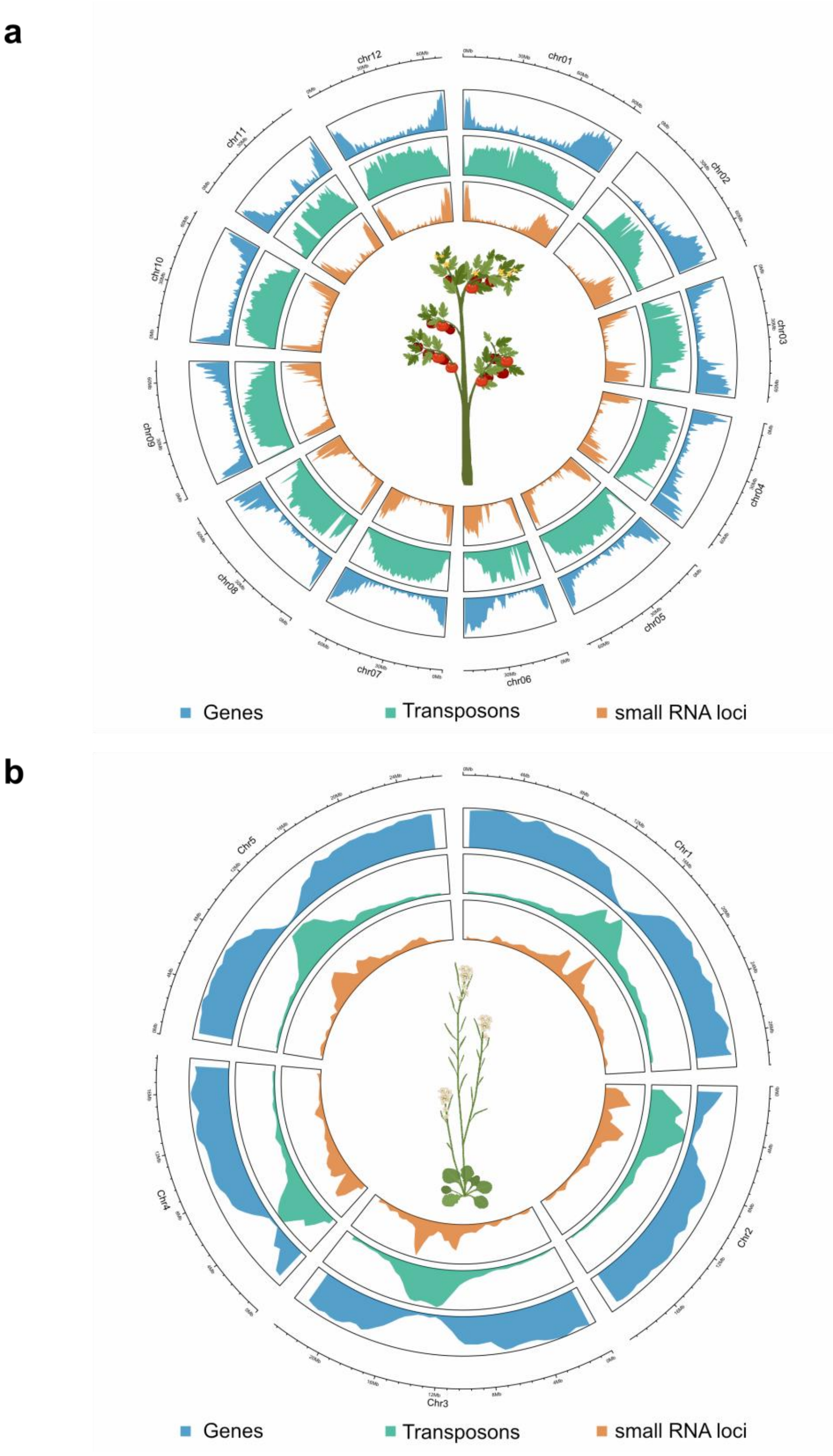


**Supplementary Fig.5. Genome-wide distribution of genes, transposable elements, and ovule small RNAs** **a.** *Solanum lycopersicum* and **b.** *Arabidopsis thaliana*. Gene density, TE density, and small RNA abundance were calculated using 1Mb genomic windows. In *A. thaliana*, ovule small RNAs are predominantly enriched in pericentromeric TE-rich regions, whereas in *S. lycopersicum*, small RNAs are mainly distributed along chromosome arms and overlap with gene-rich regions.

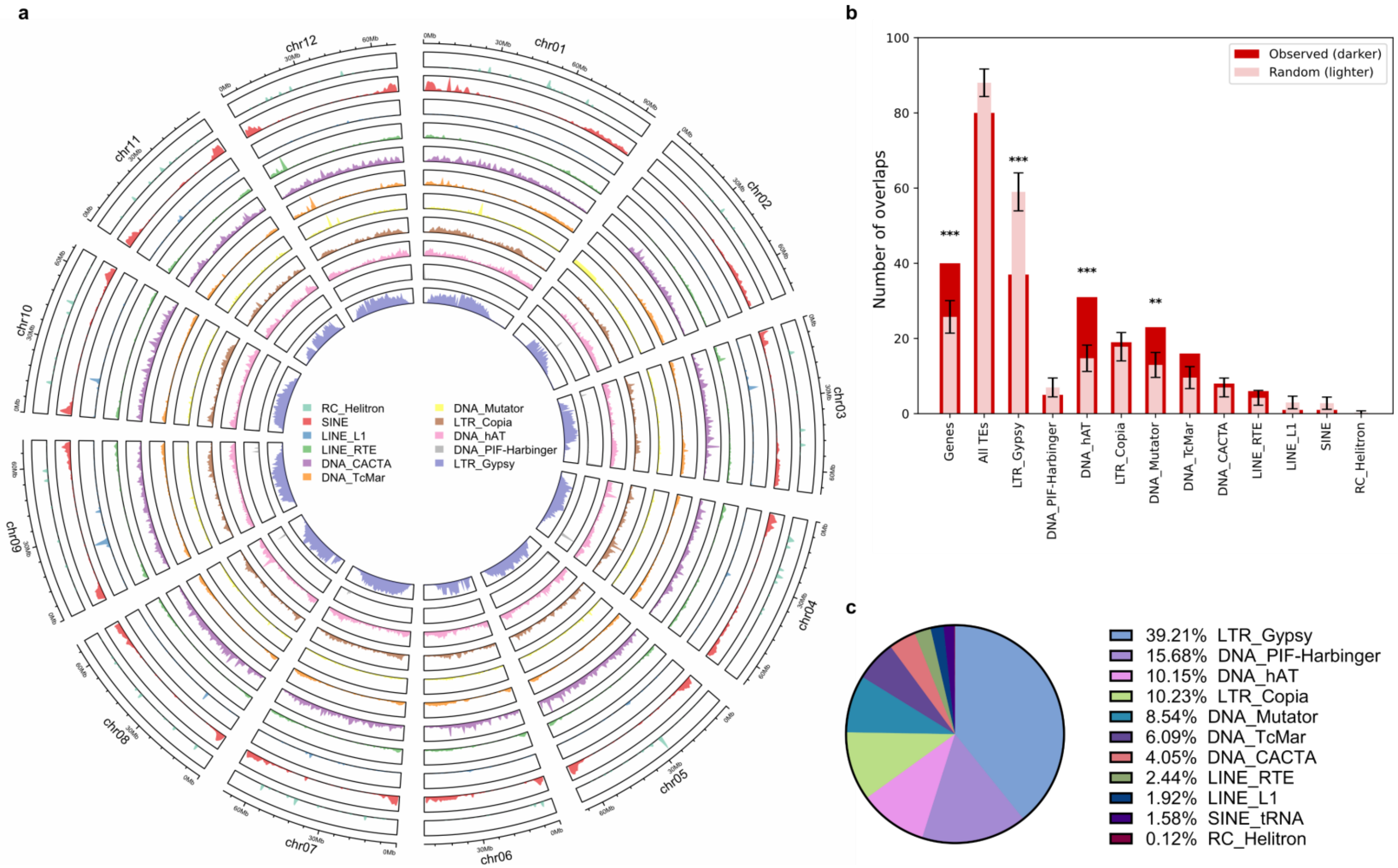


**Supplementary Fig.6. Genome-wide distribution of transposable element superfamilies in *Solanum lycopersicum*. a.** Density distribution of TE superfamilies across the *Solanum lycopersicum* genome calculated using 1 Mb genomic windows. TE superfamilies, including RC/Helitron, SINE, LINE/L1, LINE/RTE, DNA/CACTA, DNA/TcMar, DNA/Mutator, LTR/Copia, DNA/hAT, DNA/PIF-Harbinger, and LTR/Gypsy are displayed using unique colors across chromosomes. **b.** Overlap between siren loci and genomic features in *Solanum lycopersicum*. **c.** Relative abundance of TE superfamilies in the *Solanum lycopersicum* genome.

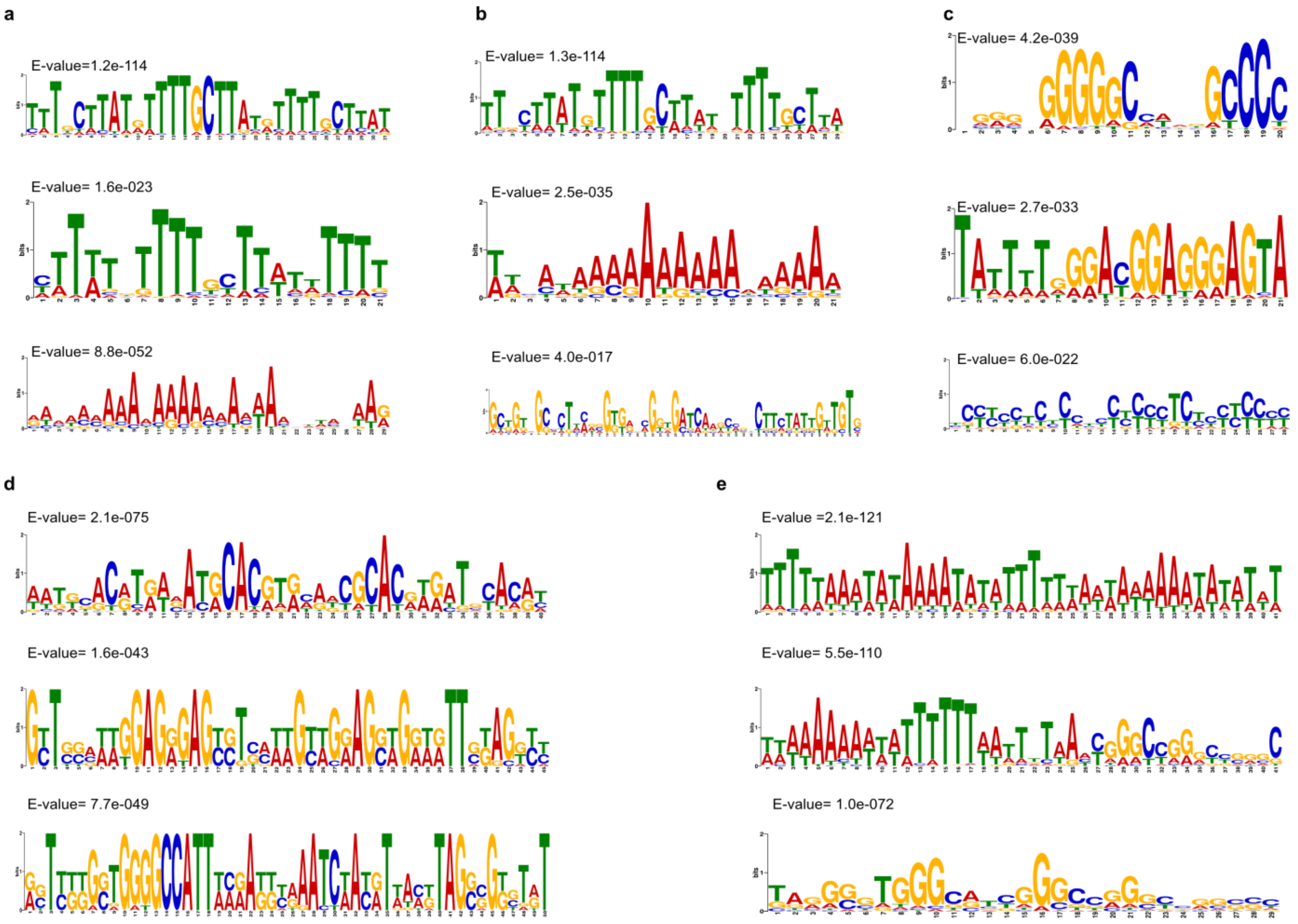


**Supplementary Figure 7. Motif enrichment analysis of ovule siren loci across species.** Motif enrichment analysis of ovule siren loci identified using MEME in **a.** *Arabidopsis thaliana*, **b.** *Capsella rubella*, **c.** *Oryza sativa*, **d.** *Solanum lycopersicum*, and **e.** *Nymphaea colorata*. The top three enriched motifs for each species are shown, with corresponding E-values indicated in the upper-left corner of each panel.

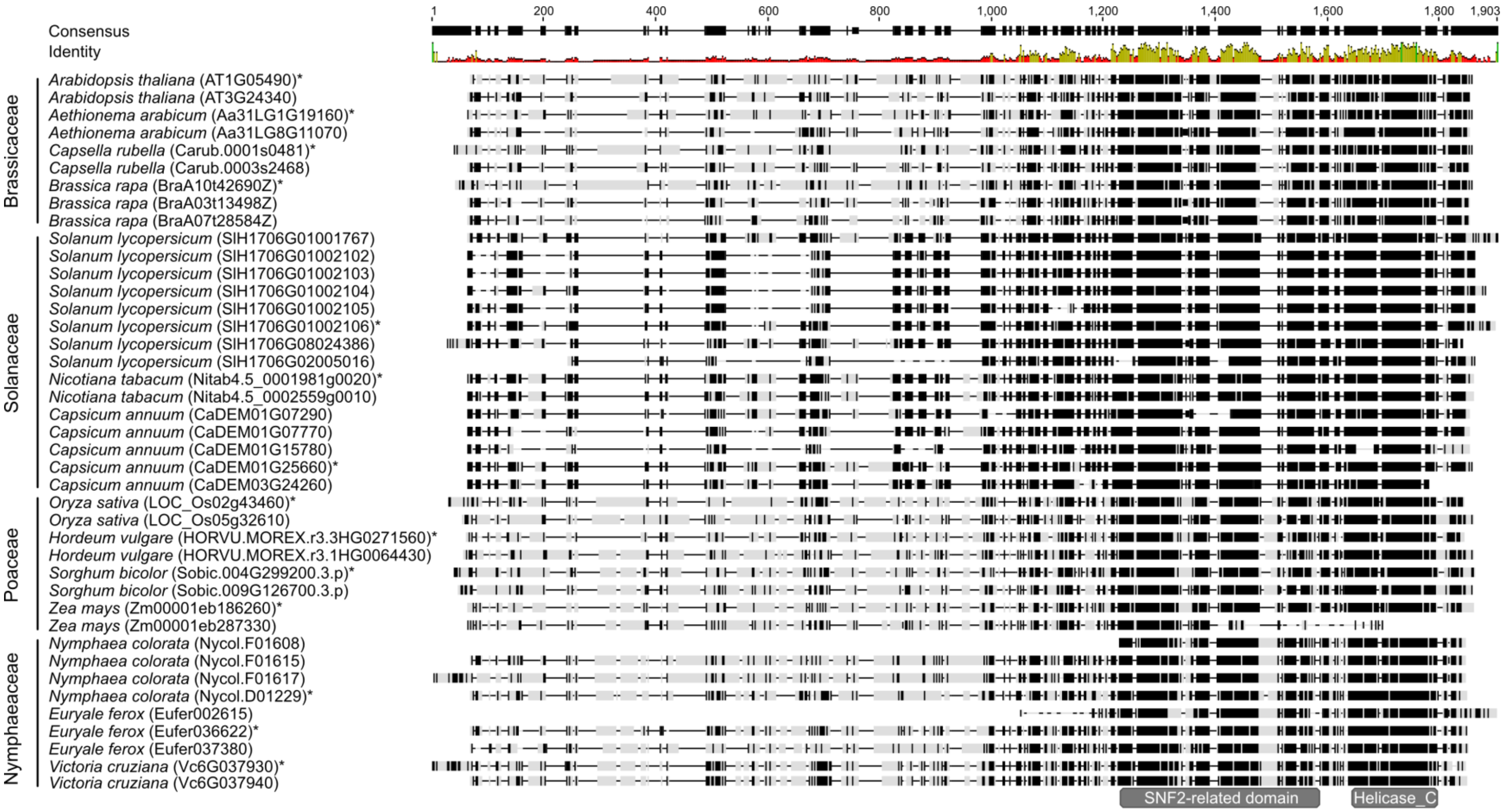


**Supplementary Figure 8. Multiple sequence alignment of CLSY3/4 proteins from representative plant species.** The alignment generated in Geneious Prime using the multiple alignment workflow with amino acid conservation visualization. Conserved SNF2 and Helicase-C domains are highlighted based on sequence similarity across species, whereas the N-terminal regions exhibit substantial divergence. Residues shaded in black indicate positions with complete similarity based on the BLOSUM62 similarity matrix implemented in Geneious Prime. Gene IDs marked with an asterisk indicate the sequences used in Fig. 4p.

60
